# Unmapped reads from whole-genome sequencing data reveal pathogen diversity in European and African cattle breeds

**DOI:** 10.1101/2025.04.17.649267

**Authors:** Daniil Ruvinskiy, Kisun Pokharel, Rodney Okwasiimire, Rayner Gonzales-Prendez, Catarina Ginja, Nasser Ghanem, Donald R. Kugonza, Mahlako L. Makgahlela, Heli Lindberg, Melak Weldenegodguad, Juha Kantanen, Martijn Derks, Richard P.M.A Crooijmans

**Author notes:** Correspondence Kisun Pokharel, Richard P.M.A Crooijmans.

## Abstract

Climate change is impacting the global spread of infectious diseases, altering pathogen distribution and transmission, threatening human and animal health. This study investigates the presence of potential pathogens in blood within unmapped reads obtained from whole-genome sequencing data of various cattle breeds across geographically diverse regions, including South Africa, Uganda, Egypt, Portugal, The Netherlands, and Finland. Unmapped reads were extracted, assembled into contigs, and subjected to taxonomic analysis based on an extensive literature search. The analysis revealed significant geographic variation in pathogen composition, with breeds in the Southern Hemisphere (Uganda, Egypt, and South Africa) showing higher alignment pathogen counts while northern breeds (particularly from Finland) exhibited lower diversity and counts. Portugal, representing a transition zone, exhibited a higher burden of parasites and tick-borne related pathogens which were also prevalent in Southern Hemisphere breeds such as *Theileria parva*, *Anaplasma platys*, *Theileria orientalis*, and *Babesia bigemina*, which is in line with the known capacity of these breeds to cope with local pathogens. Dutch breeds were found to harbor *Escherichia coli O157*, a known public health concern. The study provided key insights into emerging disease risks influenced by climate change and livestock management practices. This study highlights the potential for climate-driven variations in disease ecology and transmission, emphasizing the need for integrating genomic and environmental data and is currently the most comprehensive study to date investigating the microbial diversity present in unmapped reads obtained from WGS data of cattle populations.

**Highlights:**

1. Unmapped sequence reads’ analysis of blood reveals signatures of disease occurrence over time.
2. Blood pathogens prevail in the Southern hemisphere, becoming less evident towards northern regions (i.e. we observed a gradient pattern), with Portugal (and partly the Netherlands) showing intermediate values.
3. The commercial Holstein cattle in the six countries exhibited lower pathogen sequence alignments than their native counterparts (i.e. the Netherlands).

## Introduction

Livestock production is a cornerstone of the agricultural sector in many parts of the world, providing essential resources such as meat, milk, and fiber, while also contributing to livelihoods, particularly in rural communities. This is especially true in regions such as sub-Saharan Africa, where livestock plays a critical role in both local economies and food security (Thornton et al., 2009). However, cattle are susceptible to a range of infectious diseases, many of which are zoonotic, meaning they can also infect humans. The risk of zoonotic diseases is expected to increase in the context of climate change, with important implications for public health (Shafique et al., 2024). Changes in temperature, precipitation, and habitat suitability directly influence the ecology and behavior of vectors, reservoirs, and pathogens, leading to shifts in disease prevalence, distribution, and seasonality (Kantanen et al., 2015; McMichael, 2013; Rogers and Randolph, 2006). For instance, vector-borne diseases, including those caused by ticks, are especially sensitive to climatic conditions. As temperatures rise, the geographic range of these vectors is expected to expand, increasing the risk of disease transmission in new areas (Patz et al., 2005). Consequently, infectious diseases can have negative effects on livestock productivity, leading to reduced yields, trade restrictions, and increased veterinary costs (Nardone et al., 2006). The ongoing changes in global climate patterns are expected to exacerbate these challenges by altering the distribution, abundance, and transmission dynamics of pathogens.

Vector-borne and pathogen-induced diseases in cattle are a major constraint on livestock production across Africa. These diseases can lead to reduced milk and meat production, Increased mortality and morbidity, elevated costs due to veterinary care (e.g., tick control), vaccination programs, as well as trade restrictions and compromised food security (Knight-Jones and Rushton, 2013).

Similarly, in Europe, cattle farming is a crucial component of agricultural systems. For instance, cattle production not only supports rural economies but also plays an essential role in the conservation of certain breeds that are adapted to local environmental conditions. Breeds such as the Barrosã are well-adapted to the mountainous terrain and semi-extensive farming systems practiced in northern Portugal. These breeds are integral to the rural economy, providing high-quality meat products and maintaining traditional grazing practices that contribute to biodiversity conservation. However, like many other regions, Southern Europe is expected to experience more pronounced temperature increases, leading to longer periods of vector activity and an expansion of suitable habitats for disease vectors (Semenza and Suk, 2018). Commercial breeds of cattle such as the Holstein are more susceptible to non-native environmental conditions and local pathogens (Nardone et al., 2006).

With the advent of next generation sequencing (NGS) technologies, it is now possible to investigate the microbial diversity present in biological samples. NGS allows researchers to sequence vast amounts of genetic material from environmental or biological samples, providing insights into the presence of pathogens, even those that are not well represented in traditional reference databases (Satam et al., 2023). One of the aspects of NGS is its resultant “unmapped reads”-those sequences that do not align to the reference genome being used (i.e. to the target species), meaning that they may be representative of other organisms such as pathogens (Whitacre et al., 2015). By focusing on the analysis of unmapped reads, researchers can uncover a broader range of microbial diversity and identify potential pathogens that could be driving disease dynamics in livestock populations. This is particularly important in the context of climate change, as it allows for detection of emerging or shifting pathogen populations that may be influenced by environmental factors. Integrating genomic data with environmental and geographic information is a critical next step in understanding how climate-sensitive pathogens might spread and establish in new geographic areas (Jones et al., 2008). A few studies have explored the potential of these unmapped reads generated from WGS of blood, tissue and sperm (e.g., Neuman et al. 2023, Usman et al. 2017) to uncover viral and bacterial pathogens, providing valuable information on the microbial diversity of both environmental and biological samples. A study in Black Pied cattle found bovine parovirus 3 and Mycoplasma species (Neumann et al., 2023). Outside of cattle, unmapped DNA and RNA reads of songbirds have been analyzed to find a variety of pathogenic species including Plasmodium and Trypanosoma (Laine et al., 2019).

We focused on the analysis of unmapped reads in blood taken from the tail region and the jugular vein of a range of cattle breeds across the northern and southern hemispheres. Our aim was to investigate global pathogen trends and patterns, and the disease backgrounds of native cattle on a north-south transect. Unmapped reads were used to infer the pathogen content using a combination of custom alignment filtering and k-mer count analysis. This is a novel and very much first-of-its kind foray into the potential of unmapped reads as a method of understanding and studying diseases of cattle and the occurrence of pathogens globally.

## Materials and methods

### 2.1 Data acquisition and preprocessing

Whole-genome sequencing (WGS) data were obtained from the LEAP-Agri-project OPTIBOV on the Genetic characterization of cattle populations for optimized performance in African ecosystems (Crooijmans, 2018). OPTIBOV aimed to characterize native cattle breeds adapted to various ecosystems in Europe and Africa by sequencing the whole genomes of 27 native breeds from Europe (Finland, the Netherlands, Portugal) and Africa (Egypt, Uganda, South Africa; Figure 1). Commercial Holstein Friesian samples from each of the six countries were also included for comparison. Altogether 544 samples representing 23 breeds in 6 countries and 1 breed (HF) present in every country have been included in our study (Figure 1).

**Figure 1:**
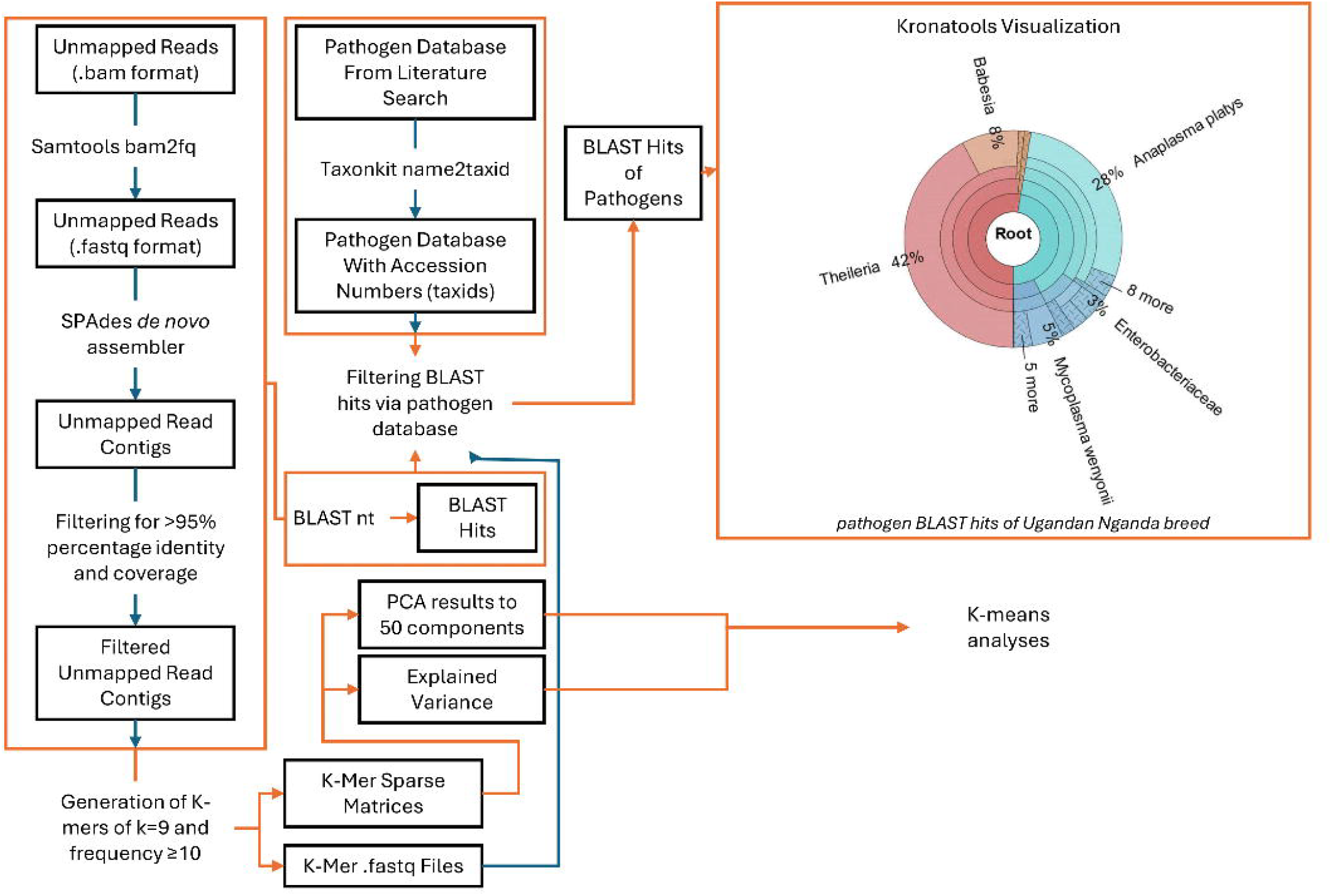
List of samples used and respective geographic locations. Samples included in the study represent 27 cattle breeds from six countries (Finland, the Netherlands, Portugal, Egypt, Uganda, and South Africa) across two continents (Europe and Africa).

The blood samples of Finnish, Dutch, South African, Egyptian, Ugandan and Portuguese cattle (including the Holstein) were collected from the jugular vein using 9 ml vaccuum collection tubes containing K3-EDTA as an anticoagulant (Greiner Bio-one reference n. 455036).

Total DNA was extracted following routine procedures common to all partners of the OPTIBOV consortium (Gonzalez-Prendes et al., 2022), and the genomic libraries were sequenced (paired-end, 150 bp) using the Illumina Novaseq 6000 platform yielding approximately 10x coverage (Gonzalez-Prendes et al., 2022). WGS data was aligned to the ARS-UCD1.2 bovine reference genome and all the sequences that did not align to the reference genome (i.e., unmapped reads) were analyzed in this study. Unmapped reads are often indicative of unknown or unexpected microbial content, including potential pathogens.

Unmapped reads were extracted from the BAM files using the Samtools (version 1.16) software (Li et al., 2009). The samtools view -f 4 command was used to extract all unmapped reads from each BAM file. These reads were subsequently converted to a FASTQ format using the samtools bam2fq command, ensuring compatibility with the downstream analysis pipeline.

### 2.2 *De novo* assembly of unmapped reads

The unmapped reads in FASTQ format were assembled into contiguous sequences (contigs) using SPAdes (version 3.13.0)(Bankevich et al., 2012), a *de novo* assembler optimized for short-read data (Bankevich et al., 2012). SPAdes employs de Bruijn graph algorithms to assemble overlapping reads into contigs, allowing us to identify novel microbial sequences not represented in the reference genome. Assembly was performed using the default parameters, and the resulting contigs were subjected to taxonomic and pathogen classification (see details below, section 2.4). Pathogens are defined by the Food and Agriculture Organization of The United Nations (FAO) as microorganisms causing disease, including viruses, bacteria, fungi, and helminths (*Foodborne antimicrobial resistance*, 2022). Contigs were further filtered to retain only sequences with >95% for both percentage identity and coverage and a minimum length of 500 bp.

### 2.3 Principal Component Analysis of unmapped reads

We performed a principal component analysis (PCA) on assembled contigs derived from unmapped reads to investigate the genetic variability across cattle breeds, . Contigs were constructed using SPAdes (Bankevich et al., 2012) and k-mer frequencies were computed with k=9. This k-mer size was chosen as it balances sensitivity and specificity for genomic sequence analysis, particularly for viral pathogen sequences. A k-mer size of 9 provides the resolution necessary to detect patterns in longer genomic regions typical of pathogens, as shorter k-mers may lack sufficient discriminatory power while larger k-mers can lead to a far too sparse dataset, reducing potential signal detection. We selected k=9 for our k-mer analysis as it represents an optimal According to Zhang et al. (2017), k=9 was found to be a suitable feature length in large-scale genomic analyses, where cumulative relative entropy (CRE) and other metrics indicated that smaller k-mer values capture sufficient diversity while maintaining computational feasibility for highly diverse datasets (Zhang et al., 2017). This value ensures robust phylogenomic comparisons while minimizing artifacts introduced by excessive feature lengths. To analyze genetic differences among breeds and countries, k-mers of size 9 were generated from unmapped reads using a sliding window approach. FASTA files containing the filtered contigs were processed to calculate k-mer frequencies for each contig. A sparse matrix representation of k-mer counts was generated using Python scripts and stored in .npz format. Low-frequency k-mers in fewer than 10 sequences, were excluded to reduce noise and computational complexity. This preprocessing step resulted in a set of k-mer matrices, one for each breed, with each matrix capturing the distribution of filtered k-mers across the contigs.4^*k*^￼ possible k-mers (262,144 for k=9) provide an effective compromise between capturing meaningful biological features and maintaining computational feasibility. K-mer frequency matrices were generated for each contig using a custom Python script employing the scipy and pandas libraries. The high dimensionality of the k-mer space necessitated dimensionality reduction through PCA, implemented using the scikit-learn library. Sparse matrices were used to optimize memory usage during the computation. The first principal components, explaining the majority of variance, were used for clustering and visualization of contigs by breed, highlighting genetic relationships and potential pathogen-driven variations among global cattle populations.

Clustering was performed to explore genetic relationships at both the country and breed levels. K-means clustering was applied to the PCA-transformed data for two scenarios: 6 clusters (countries) and 27 clusters (breeds). The silhouette score was used to determine the optimal number of principal components for each clustering scenario. For 6 clusters, the highest silhouette score was observed at 20 principal components, while for 27 clusters, the optimal number was 30 principal components. These dimensionalities were subsequently used for all clustering and visualization analyses.

PCA scatter plots were generated to visualize clustering patterns for both scenarios. For 6 clusters, PCA results using 20 components were plotted, with each point color-coded based on its assigned country cluster. Similarly, for 27 clusters, PCA results with 30 components were visualized, highlighting breed-specific clusters. To further aid interpretation, an additional PCA visualization was performed where each breed was represented by a single point, derived from the mean PCA scores of all contigs belonging to the breed. Breed names were overlaid on the plots to identify their positions within the PCA space.

### 2.4 Taxonomic analysis using KronaCharts and BLAST

The assembled contigs were subjected to taxonomic analysis to identify potential pathogens. First, BLAST (Basic Local Alignment Search Tool) searches (Altschul et al., 1990) were conducted against the non-redundant NCBI nucleotide database to identify homologous sequences. The output from BLAST was then processed using KronaTools (Ondov et al., 2011) to visualize the taxonomic composition of the sequences (Ruvinskiy, D. and Pokharel, K., 2025). KronaCharts provides interactive, hierarchical visualizations of the microbial taxa present in each sample at various levels (e.g., kingdom, phylum, genus, species), allowing for detailed exploration of pathogen diversity. BLAST results were filtered using the pathogen database generated from the literature search, and contigs matching known pathogens of interest were prioritized for further analysis. The pathogen database and NCBI TaxIDs from TaxonKit (Shen and Ren, 2021) ensured that identified pathogens were relevant to the breeds and regions studied.

### 2.5 Pathogen database and literature search

To facilitate the identification of relevant pathogens, a comprehensive pathogen database was created based on a systematic literature search in NCBI (see supplementary file 1). The search focused on pathogens known to affect cattle in the regions studied, including both endemic and emerging infectious agents. Searches were conducted across peer-reviewed literature using keywords related to cattle pathogens, specific breeds, and geographic regions. For example, for a literature search of Ugandan pathogens the keywords “Ugandan cattle pathogens, disease” were used. Articles from major veterinary, microbiology, and zoonotic disease journals were included in this database, using 12 articles in total. Each identified pathogen was cross-referenced with available NCBI Taxonomy identifiers (TaxIDs) to ensure consistency and to facilitate downstream analysis. This database was then used to filter the assembled contigs for potential pathogens of interest.

### 2.6 Taxonomic resolution using TaxonKit

To ensure that pathogen identification was accurate and up to date, we used TaxonKit v. 0.19.0 (Shen & Ren, 2020), a command-line toolkit for taxonomic data manipulation. TaxonKit was employed to map scientific names of the pathogens identified in our literature search to their respective NCBI Taxonomy identifiers (TaxIDs). TaxonKit commands such as taxonkit name2taxid were used to convert the names of identified pathogens into their corresponding TaxIDs as well as the TaxIDs of subspecies and variants of these pathogens. Additionally, where the literature did not provide a TaxID, TaxonKit’s taxonkit list function was used to retrieve and verify the correct TaxIDs from the NCBI taxonomy database. Pathogen genus were used to get taxids of all potential related species of pathogen. These identifiers were then used to match against the taxonomic output from the BLAST searches, ensuring that the identified pathogens were accurate and up to date with the latest taxonomic revisions. In total 89,785 potential pathogens were identified as a result.

### 2.7 Comparative pathogen profiling

The identified pathogens from the taxonomic and statistical analyses were further compared across regions to identify patterns of geographic variation using Python 3.10.6. Pathogen counts were correlated with environmental and climatic factors in each region to explore the potential influence of climate on pathogen diversity and prevalence. Additionally, the relative abundance of pathogens within each breed was examined to assess whether certain cattle breeds exhibited unique susceptibility to pathogens.

The above described pipeline for pathogen sequence detection in unmapped reads is presented in Figure 2.

**Figure 2:**
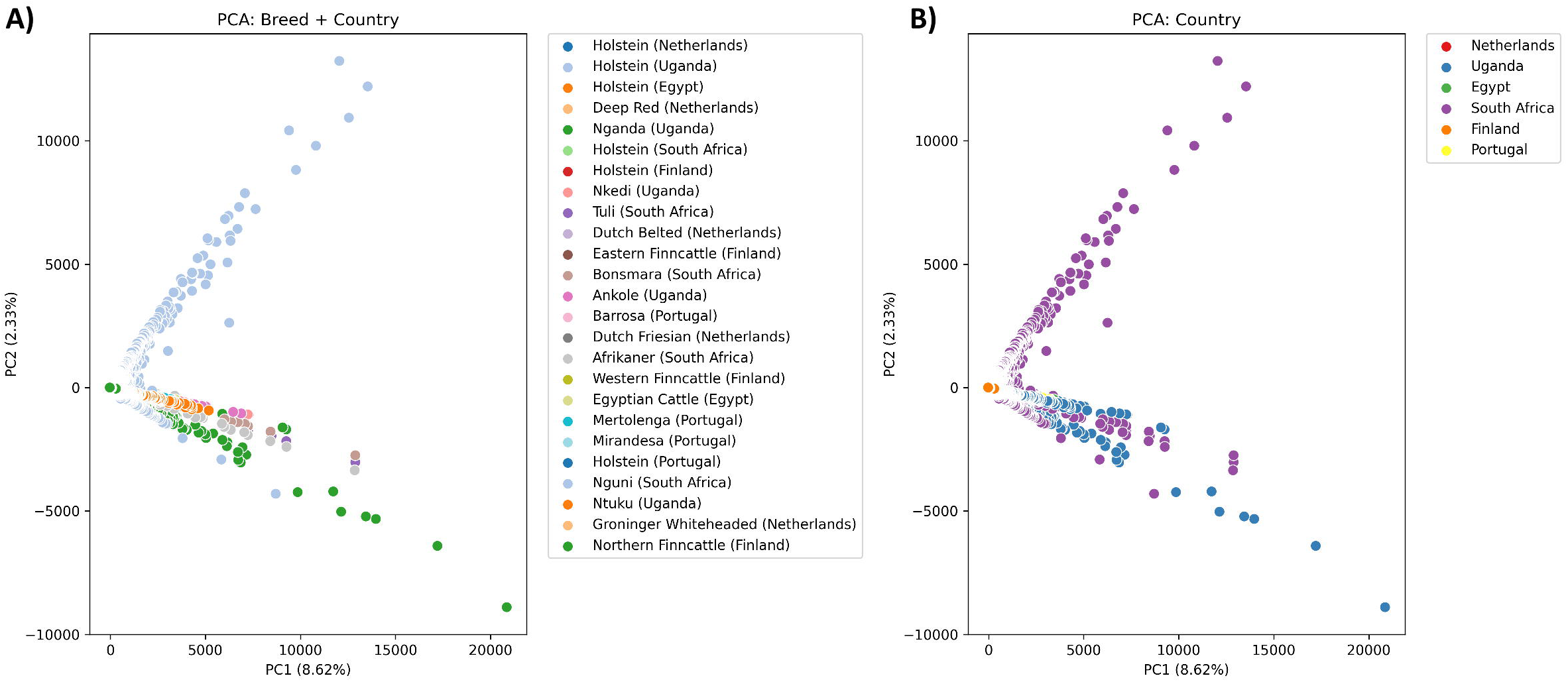
Flowchart showing the pipeline used for analyzing unmapped reads.

## 3. Results

### Results from *de novo* assembly

The assembly results were analyzed to assess the quality and characteristics of contigs before and after filtering. Prior to analysis there were 1,327,848,784 reads in all unmapped read files with an average 249,127 reads per sample. In total, SPAdes assemblies across all datasets generated 3,358,783 unfiltered contigs (supplementary file 2), with a cumulative length of 1,558,751,633 bp and an average GC content of 46.72%. The N50 length of the unfiltered assemblies ranged from 233 bp to 4,247 bp, while the largest contig reached a length of 962,072 bp, reflecting the assembly’s ability to capture large genomic regions. Filtering significantly improved the quality of the assemblies by removing shorter (< 500 bp) and lower-confidence (>95% coverage and sequence identity) contigs. After filtering, the total number of contigs was reduced to 506,868, corresponding to a cumulative length of 873,470,670 bp. The N50 length increased to a range of 740 bp to 10,532 bp, indicating improved contiguity in the filtered datasets. Additionally, the GC content remained consistent at 46.42%, further validating the accuracy of the assembly process. Among the datasets, the largest filtered contig measured 962,072 bp, and the number of high-quality contigs (≥1,000 bp) increased significantly post-filtering.

#### 3.1 Principal Component Analysis of unmapped reads

Principal Component Analysis (PCA) was performed on k-mer frequency matrices derived from assembled contigs of unmapped reads across cattle breeds. The PCA, calculated to 50 principal components, revealed that the variance was distributed across multiple components. This is consistent with the high dimensionality and complexity of a dataset of unmapped read contigs. The first principal component (PC1) explained 8.62% of the total variance, while the first 10 principal components collectively accounted for 18.19% of the variance. All 50 calculated PCs had a cumulative variance at ∼22.35%. This distribution of variance indicates a more complex set of data with no completely dominating trend. Despite this, some clustering patterns were evident when visualizing the breeds and countries in PCA space. A scatterplot of the first two principal components (PC1 and PC2) revealed both clustering and overlap among breeds (Figure 3), reflecting their genetic diversity and potential shared pathogen-related sequences. Specific breeds such as the Ugandan Nkedi formed distinct clusters, while others overlapped. This is indicative of shared genetic or environmental factors influencing pathogen presence. Most interestingly, Holstein samples appear across multiple clusters rather than forming just one group. The scattering of Holstein suggests that the unmapped k-mer profiles are influenced by geographic factors. This suggests that unmapped sequences are not simply a result of the host genome and are strongly affected by external factors that vary by geographic location.

**Figure 3:**
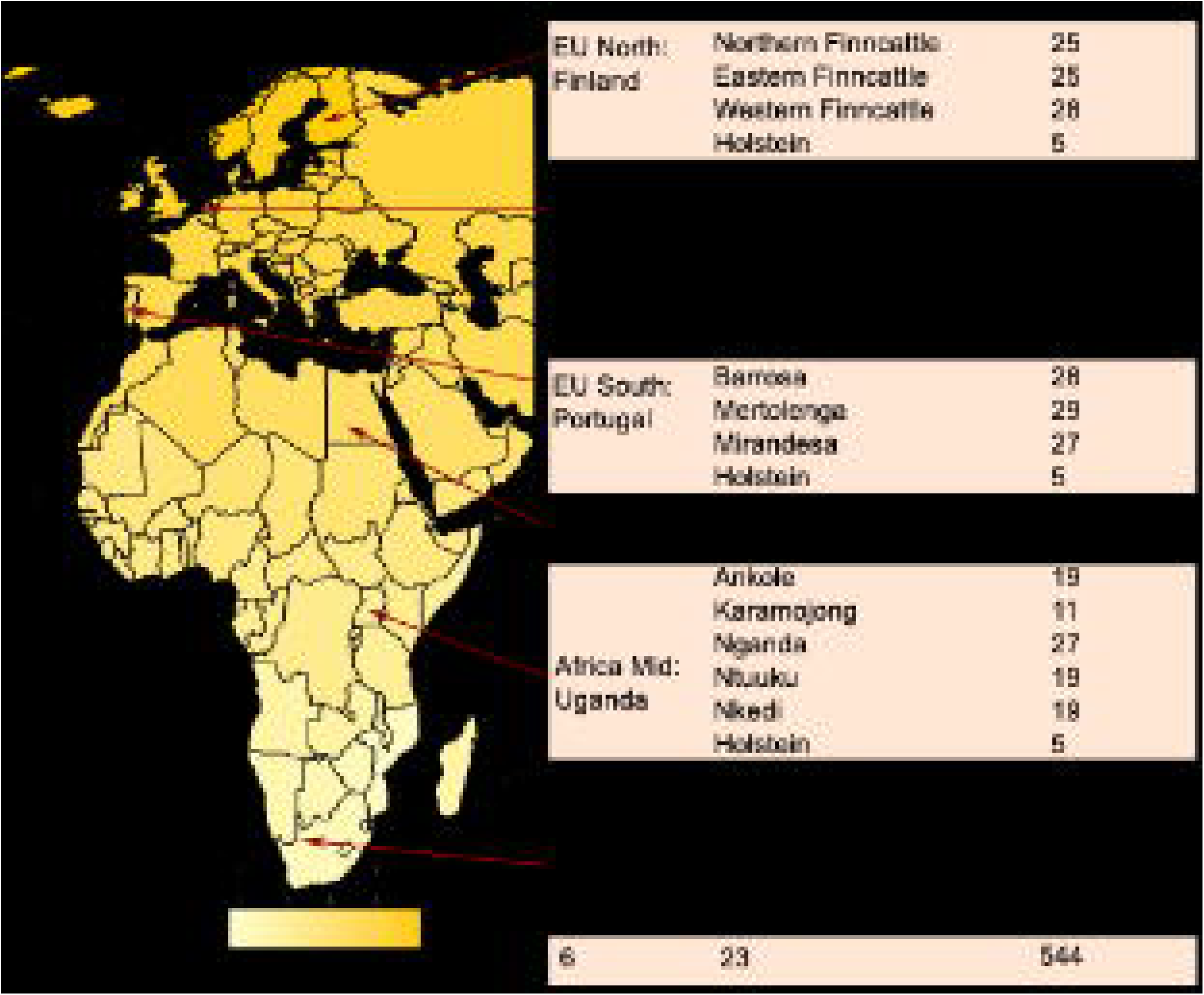
Principal Component Analysis visualization of cattle breed k-mers of size 9. Visualization A shows the clustering of breeds, whereas B shows the clustering according to country.

#### 3.1 Total pathogen counts across breeds and regions

A comparison of total pathogen alignment counts across cattle breeds (Ruvinskiy, D. and Pokharel, K., 2025) and regions revealed significant variation, indicating potentially differing levels of exposure among the studied populations. A variety of studies from respective countries in the northern and southern hemispheres were used to build a database for filtering pathogens (supplementary file 1). In total 89,785 potential pathogens were identified.

The total number of pathogens related sequences detected varied significantly across regions, with some regions showing much more alignments to pathogen related sequences compared to others. Uganda had the highest total pathogen count, with 527 pathogen related sequences identified across Ugandan cattle breeds (Table 1). South Africa, the Netherlands and Portugal also exhibited high pathogen counts, with 396, 236 and 119 pathogens detected, respectively. Finland and Egypt reported 59 and 58 total pathogen related sequences (Table 1, supplementary file 3).

**Table 1:**
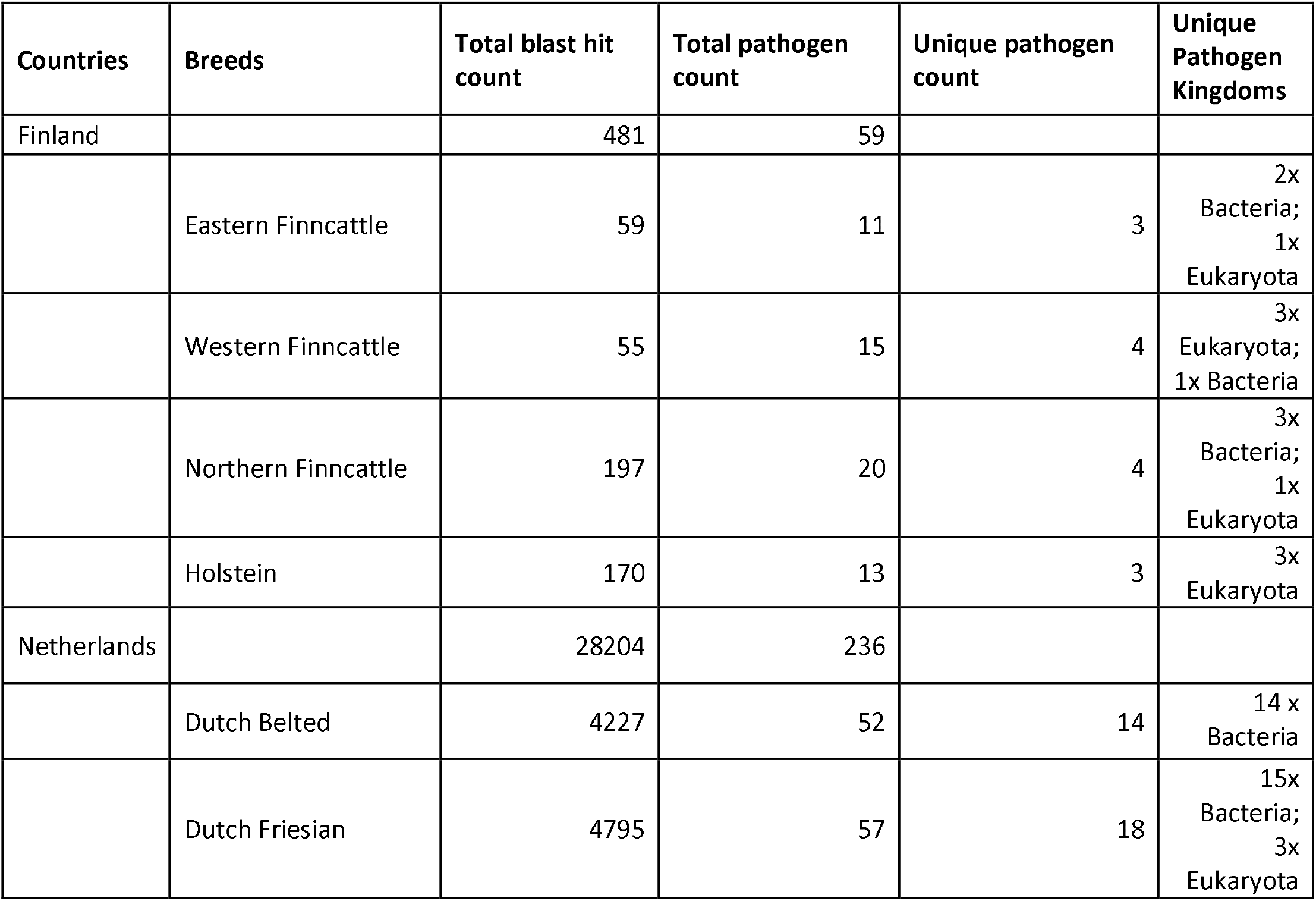
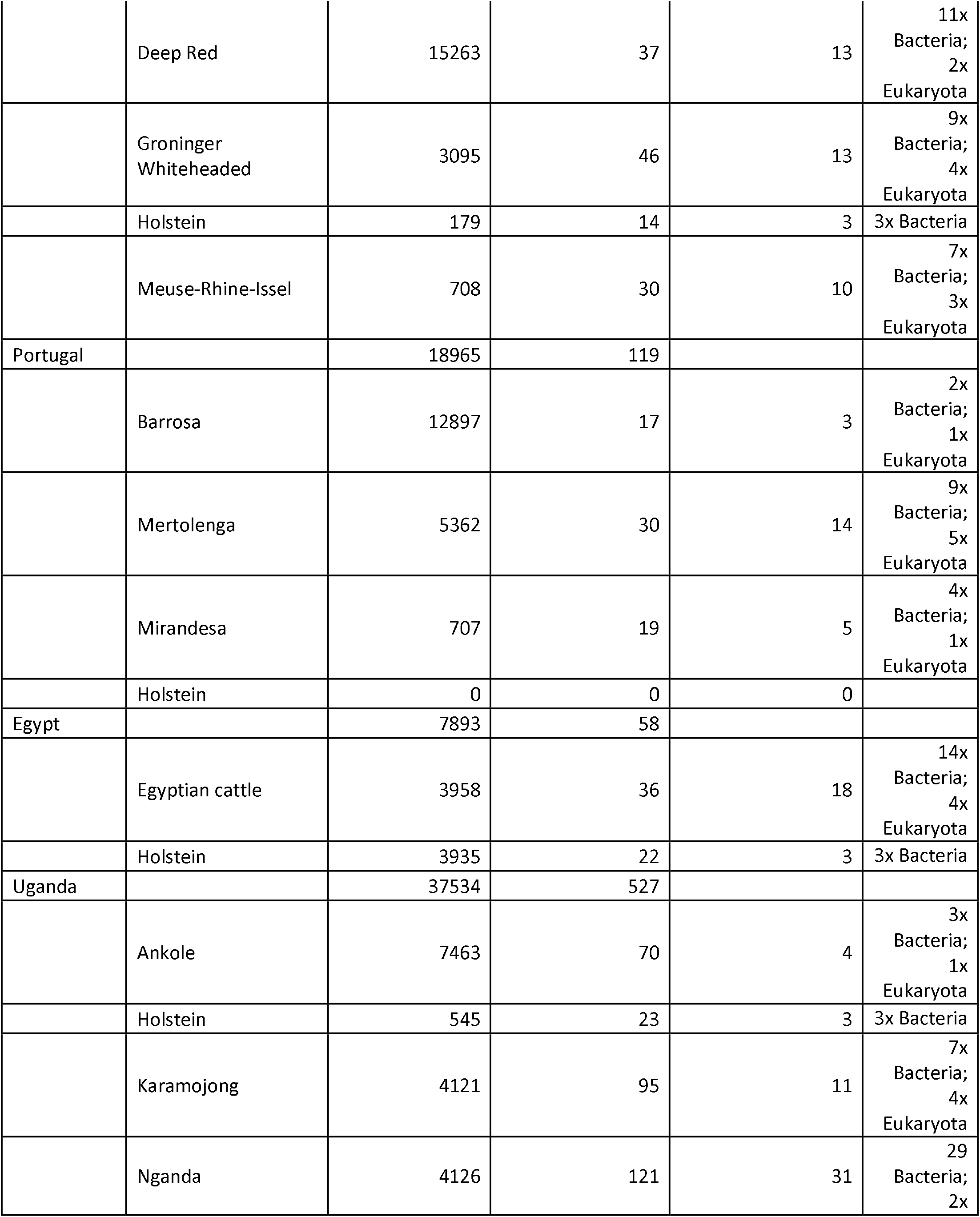
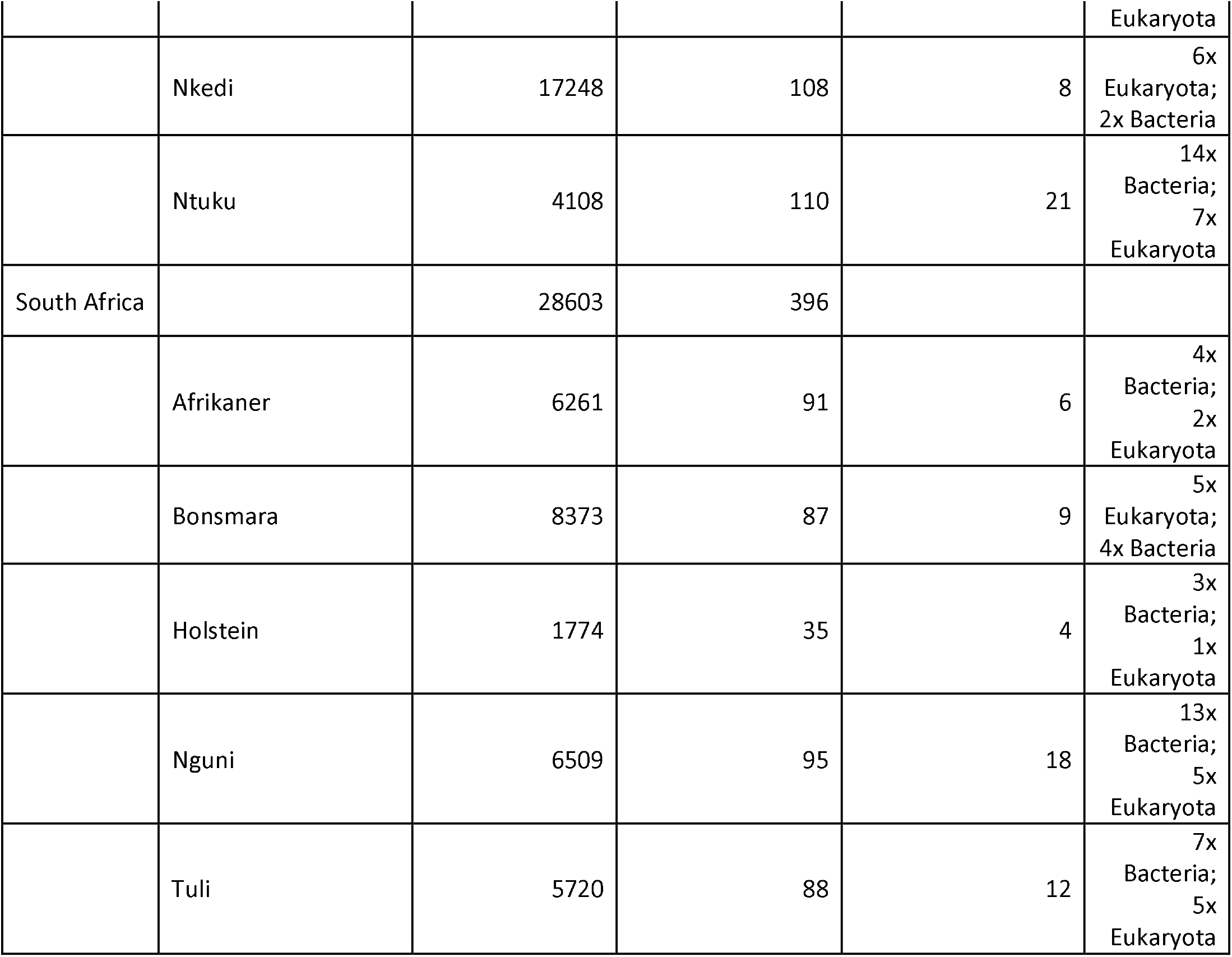
Total number of blast hits post-filtering, and total and unique pathogen taxa counts for countries and breeds.

At the breed level, several breeds showed significant pathogen counts, particularly in regions where environmental conditions are favourable for pathogen survival. The Nganda cattle from Uganda had the highest total pathogen count, with 121 pathogens detected (Table 1), suggesting that these cattle are exposed to a broad range of pathogens. In contrast, the Eastern Finncattle from Finland exhibited the lowest pathogen count, with only 11 pathogens detected (Table 1). Dutch Belted cattle from the Netherlands showed a moderate pathogen sequence presence, with 52 pathogens detected (Table 1), potentially reflecting the temperate climate in this region rather than controlled farming practices as it is a year-round grazing breed.

Overall, the breeds from southern regions such as South Africa and Uganda showed higher pathogen loads, likely due to environmental conditions that favor pathogen proliferation and transmission, while breeds from temperate regions exhibited lower pathogen counts. The absence of pathogens in Portuguese Holsteins is particularly notable. This could reflect enhanced biosecurity measures or potential sampling or sequencing biases.

#### 3.3 Pathogen diversity across breeds and regions

A wide variety of pathogens were identified across the cattle breeds studied, primarily consisting of bacteria and a few eukaryotic pathogens. Table 1 provides a summary of the unique pathogen counts for each breed, reflecting the diversity of pathogens identified in all samples in a breed.

In Uganda, indigenous breeds such as Nkedi (108 unique pathogens) and Nganda (121 unique pathogens) demonstrated the highest pathogen diversity. The imported Holstein breed, commonly used for dairy production, showed a much lower pathogen diversity, with only 3 unique pathogens identified.

Supplementary file 2 provides an overview of the unique pathogens identified across different cattle breeds and countries. Each breed is associated with a distinct set of pathogens, reflecting regional variations in pathogen exposure.

#### 3.4 Pathogen composition across breeds and regions

##### Pathogen profiles in the Finnish Cattle Breeds

There are three native breeds in Finland: Western Finncattle, Northern Finncattle, and Eastern Finncattle. These breeds are adapted to the cold climate and are typically raised in small scale farming systems. While vector-borne diseases are less prevalent in colder climates, there is concern that rising temperatures may allow new pathogens to emerge in northern Europe (Patz et al., 2005). Among the four Finnish cattle breeds—Eastern Finncattle, Holstein, Northern Finncattle, and Western Finncattle— several pathogens were found to be common across all breeds. Notably, the following pathogens were identified in all four breeds: *Candidatus Mycoplasma haemominutum ‘Birmingham 1’*, *Mycoplasma haemocanis str. Illinois*, *Mycoplasma haemofelis Ohio2*, *Mycoplasma ovis str. Michigan*, and *Mycoplasma parvum str. Indiana*. These common pathogens indicate a shared exposure across Finnish breeds, which may be reflective of environmental factors or common management practices in Finnish cattle farming. The presence of these specific Mycoplasma species across all breeds suggests that these pathogens might be prevalent in the Finnish cattle population and could be a focus for disease management programs. In addition to common pathogens, each Finnish breed also exhibited a distinct set of unique pathogens, indicating breed-specific pathogen-host interactions. Eastern Finncattle was uniquely associated with pathogens such as *Eimeria maxima* and *Mycoplasma haemocanis*. These pathogens suggest exposure to parasitic infections, with *Eimeria maxima* being particularly associated with intestinal coccidiosis in avian species(Jebessa et al., 2022). Northern Finncattle exhibited unique pathogens including *Candidatus Mycoplasma haemobos*. The presence of *Candidatus Mycoplasma haemobos* is interesting as it has previously been confirmed in Korean and German cattle (Hoelzle et al., 2011; Kim et al., 2024). Western Finncattle showed pathogens such as *Alentia gelatinosa* and *Eimeria tenella*, the latter being a well-known cause of coccidiosis in broilers (Choi et al., 2021). This finding suggests a potential higher risk of parasitic infections in this breed.

##### Pathogen Profiles in the Dutch Cattle Breeds

The Netherlands is home to several dual-purpose breeds but selected for dairy purpose, including the Groningen White Headed and Dutch Frisian breeds. The Deep Red and Dutch Belted are more selected as meat type animals. Dairy type traditional cattle are typically raised in more intensive farming systems, where biosecurity measures are critical to controlling the spread of infectious diseases, while the Dutch belted breed is outside year-round. However, the increasing occurrence of extreme weather events poses new risks for Dutch cattle populations by potentially altering the distribution of disease vectors (McMichael, 2013). From the analysis of unmapped reads across multiple breeds of Dutch cattle, several pathogens were identified as common across all Dutch breeds. These pathogens include *Anaplasma phagocytophilum, Pseudomonas aeruginosa, and Escherichia coli*. The presence of these pathogens suggests a widespread occurrence in the Dutch cattle population, indicating potential endemicity or common exposure to similar environmental or management conditions.

In addition to the common pathogens, there were also those found to be unique to specific breeds. In the Dutch Belted cattle, a variety of *Salmonella enterica* subspecies were aligned, including *Salmonella enterica subsp. enterica* serovar *Newport*, *Bareilly*, *Kentucky* and *Stanley*. A number of *E. Coli* strains were also detected including *O18*, *O157*, *O1*, and *O99*. A similar profile can be observed in the Deep Red, Holstein and the Friesian with the latter being observed to also align with *Klebsiella oxytoca* and *Pseudomonas aeruginosa*-gram negative bacterium known to cause mastitis (Massé et al., 2020). In the Meuse-Rhines-Yssel *Candidatus Mycoplasma haemobos* was also located. This pathogen infects the red blood cells of cattle (De Souza Ferreira and Ruegg, 2024). In addition *Spirometra erinaceieuropaei* and *Mycoplasma yeatsii* were also located in this breed. The former is a tapeworm typically infecting carnivore and amphibian hosts as well as humans, but cattle does not seem to have been identified as a reservoir thus far; the latter is a mycoplasma that typically affects goats (Calcutt et al., 2015), but has previously been located in cattle ear canals (Amadou Sery et al., 2024).

##### Pathogen Profiles in the Portuguese Cattle Breeds

Portuguese cattle breeds, including Barrosã, Mertolenga, and Mirandesa, are integral to the country’s rural economy. The country, of course, also boasts a commercial Hostein population. The Barrosa breed is a dual-purpose breed, valued for both its meat and milk production. It is raised in the mountainous regions of northern Portugal, where semi-extensive grazing systems dominate. The potential impact of climate-sensitive pathogens, particularly those spread by ticks, is an emerging threat to these cattle populations, as southern Europe is expected to face rising temperatures and increased vector activity in the coming decades (Semenza and Suk, 2018). The analysis of pathogen profiles in the Portuguese cattle breeds—Barrosa, Mertolenga Mirandesa, and Portuguese Holstein—revealed both common and unique pathogen occurrences among the breeds. The Portuguese Holstein breed, notably, did not show any alignments to pathogens in the BLAST analysis. This absence could be indicative of several factors: potentially lower pathogen exposure in the commercial environment or technical factors such as differences in sample collection or sequencing depth. This lack of detectable pathogens in the Portuguese Holstein presents an interesting contrast to the other Portuguese breeds, which exhibited a variety of unique pathogens. The remaining breeds had some pathogens in common: *Theleria orientalis, Anaplasma marginale* and *ovis*, as well as a range of *Mycoplasma* species. *Theleria orientalis* has already been characterized in Portuguese cattle, specifically in the Mirandesa (Brigido et al., 2004). In addition, this pathogen has been detected in asymptomatic cattle as well (Gomes et al., 2013). *Anaplasma marginale* has been detected in *Rhipicephalus bursa* ticks in Portugal (Ferrolho et al., 2016) and *Anaplasma ovis* has been detected in wildlife vectors, sheep and cattle in other areas of Europe. In the Barrosa breed, three unique pathogens were identified, including *Anaplasma phagocytophilum* strains (*Norway variant 1* and *variant 2*) and *Schistosoma mattheei*. The Mertolenga breed demonstrated a broader range of unique pathogens, including B*abesia bigemina, Candidatus Mycoplasma haemolamae str. Purdue, and Harmothoe impar*. Similarly, the Mirandesa breed showed unique pathogen profiles, including *Anaplasma marginale (Gypsy Plains and St. Maries* strains*), Escherichia coli, and Theileria equi*. These findings indicate a clear diversity in pathogen exposure among the different Portuguese breeds, with Mertolenga and Mirandesa showing the highest number of unique pathogens at 14 and 5 (supplementary file 2). The absence of pathogens in the Portuguese Holstein breed provides a significant point of contrast and suggests potential differences in pathogen resistance or environmental exposure. The analysis of pathogen profiles among the Portuguese cattle breeds—Barrosa, Mertolenga, and Mirandesa—revealed several common pathogens shared between at least two of the breeds. These common pathogens include: *Anaplasma ovis str. Haibei, Anaplasma phagocytophilum str. Dog2*, *Schistosoma curassoni*, *Anaplasma marginale str. Florida*, and *Anaplasma centrale str. Israel*. The presence of *Anaplasma* species across these breeds suggests a significant exposure to tick-borne diseases, as *Anaplasma* is commonly associated with diseases like anaplasmosis in cattle (Santos et al., 2009). The detection of *Schistosoma curassoni*, a parasitic flatworm, indicates potential exposure to schistosomiasis, a disease typically associated with tropical and subtropical regions(Léger et al., 2016; Namirembe et al., 2024). The shared presence of these pathogens across multiple breeds underscores the common environmental and epidemiological pressures faced by these cattle breeds in Portugal.

##### Pathogen Profiles in the Egyptian Cattle Breeds

Egypt’s cattle populations, including Egyptian cattle and Holstein, are primarily raised in intensive farming systems along the Nile River and its delta. Egypt’s hot and arid climate poses unique challenges for livestock management, with diseases such as Pasteurella infections and Anaplasmosis being of particular concern. The increasing frequency of extreme weather events, such as heatwaves, is likely to exacerbate these challenges by increasing pathogen load and vector populations (Patz et al., 2005). Egyptian cattle and Egyptian Holstein revealed both shared and unique pathogens across the two groups. The Egyptian cattle breed exhibited a wider diversity of pathogens compared to the Egyptian Holstein, with a total of 3 unique pathogens identified. Among these, notable pathogens included *Anaplasma marginale, Babesia bigemina, Listeria grayi, Mycoplasma wenyonii, parvum* and *ovis* and *Schistosoma margrebowiei*. These pathogens are associated with various cattle diseases such as anaplasmosis, babesiosis, and listeriosis, which may have significant impacts on the health and productivity. *Babesia bigemina* in particular is a serious problem in Egypt and is considered one of the most worrying endemic parasitic diseases affecting cattle there (Adham et al., 2009; Mahmoud et al., 2024). In contrast, the Egyptian Holstein breed showed a more restricted pathogen profile, with only three unique pathogens: *Anaplasma phagocytophilum str. Norway variant1, Pasteurella multocida*, and *Ehrlichia ruminantium str. Welgevonden. Pasteurella multocida* is known to be a causative agent of respiratory diseases in cattle (Namba et al., 2025), while *Anaplasma phagocytophilum* variants are associated with tick-borne diseases (Apaa et al., 2023). Both breeds shared several common pathogens, including *Theileria annulata, Anaplasma marginale* (with multiple strains), *Pseudomonas aeruginosa, Mycoplasma wenyonii*, and *Theileria parva*. These common pathogens suggest a similar exposure to endemic diseases in the Egyptian environment.

##### Pathogen Profiles in the Ugandan Cattle Breeds

Uganda’s cattle industry is vital, with breeds such as Karamojong, Ntuku, and Ankole forming the backbone of rural livelihoods. Uganda’s climate, characterized by tropical conditions and seasonal rainfall, creates an ideal environment for the spread of vector-borne diseases such as East Coast fever, caused by *Theileria parva*. As temperatures continue to rise in East Africa, the risk of these diseases spreading to new areas is a growing concern (Thornton et al., 2009). The analysis of pathogen presence across Ugandan cattle breeds revealed both similarities and differences between their pathogen profiles.

Among the Ugandan breeds analyzed, seven pathogens were shared: *Babesia bigemina, Escherichia coli, Mycoplasma ovis (str. Michigan), Mycoplasma wenyonii (str. Massachusetts), Pseudomonas aeruginosa, Pseudomonas aeruginosa PA96*, and *Theileria parva (str. Muguga)*. These pathogens, shown to be prevalent across all breeds, indicate a pervasive risk of tick-borne diseases and other bacterial infections in Uganda’s cattle populations. All these breeds, regardless of management or location, are exposed to similar pathogen pressures which tracks with the real-world situation. Four of these common pathogens are vector diseases-*Babesia* and *Theleria* are transmitted by ticks (*Rhipicephalus boophilus micropus* and *Rhipicephalus appendiculatus* respectively) (Mahmoud et al., 2024); *Mycoplasma wenyonii* and *ovis* are both transmitted through various blood sucking insects, biting flies as well as contaminated equipment (Amadou Sery et al., 2024; Kim et al., 2024). What is interesting about the aforementioned *T*.*parva Muguga* strain is that it is often times used in a vaccine dubbed “the Muguga cocktail”, at times responsible for carrier status in individuals (Oligo et al., 2023). Having both *Pseudomonas aeruginosa* and *Pseudomonas aeruginosa PA96* is an interesting addition here. As well as exhibiting the more widely known strain of the pathogen, the common pathogen profile also exhibits *PA96*, which is multidrug resistant-resistant strains have been prevalent in Uganda at a household and farm level, but the latest research as of 2023 is yet to embark on multi-locus sequence typing to provide more precise information on the identity amongst different strains (Badawy et al., 2023).

The analysis of Ugandan cattle breeds revealed distinct pathogen profiles across the various breeds as well. The Ankole breed was found to host *Brucella sp. 09RB8471, Theileria parva* glutamine rich membrane protein mRNA, *Mycobacterium avium paratuberculosis* and *Brucella melitensis. Brucella* species are zoonotic, with *B. melitensis* being the most common species of brucella in human illnesses and typically associated with sheep and goats as a specific animal host, and more rarely in camels and cattle in some regions with extensive small ruminant populations (Diaz Aparicio D., 2013). Ugandan cattle breeds have been known to be infected with *B. melitensis* (Miller et al., 2016). *Theileria parva parva* glutamine rich membrane protein mRNA presence could be due simply to the BLAST database annotating a genomic region to a known transcript due to a close representation to coding exon regions. *Mycobacterium avium* is has also been described in Uganda cattle and has been a source of concern (Okuni et al., 2012). In the Holstein breed, unique pathogens included *Candidatus Mycoplasma haemobos*, which has been located in Ugandan cattle and can develop clinical signs including anemia, transient fever, lymphadenopathy and anorexia, though in most cases remaining subclinical (Byamukama et al., 2020). The Karamojong breed exhibited a diverse array of unique pathogens, including *Candidatus Brucella* strains, *Trichobilharzia regenti, and Anaplasma marginale str. South Idaho, Salmonella and Mycobacterium bovis* variant *bovis BCG* strain. These pathogens are associated with diseases like anaplasmosis and babesiosis, which can cause significant morbidity in cattle due to parasitic and bacterial infections (Fesseha et al., 2022). For the Nganda breed, unique pathogens such as *Salmonella enterica, Escherichia coli O157:H7, Brucella sp. MAB-22*, and a Variety of *Proteus* species were identified. These pathogens suggest the potential for exposure to both tick-borne diseases and bacterial infections in this breed. In the Nkedi breed, notable pathogens included *Trypanosoma vivax, brucei and cruzi, Prototheca wickerhamii, Mycoplasma haemocanis*. These pathogens are associated with trypanosomiasis, a vector-borne disease transmitted by tsetse flies (a pest which covers 70-75% of Uganda’s landmass), and protothecosis, which can affect both animals and humans (Ramadán et al., 2025). Indeed, the Nkedi breed is rarely sprayed with acaricide as the animals are rarely found with ticks. It was assumed that the results would show significant absence of tick born pathogens and indeed, that appears to be the case. The Ntuku breed showed a unique profile with pathogens such as *Trypanosoma equiperdum, Schistosoma mattheei, Escherichia coli O1:H42 Escherichia coli O125ac:K+:H10, Escherichia coli O15:H12*, and *Salmonella enterica*. The presence of *Schistosoma* is interesting here as this has been a problem for Ugandan cattle, especially in the West of the country (Namirembe et al., 2024). It is important to note that Nganda and Nkedi are kept in areas with heavy human population density so a build up of pathogens is more likely than the other breeds. The individuals here are also are from near the Lake victoria basin with more rainfall and humid conditions, while the other 3 breeds (Ankole, Ntuku and Karamojong) are from much drier zones.

##### Pathogen profiles in South African cattle breeds

South Africa is home to several indigenous and commercial cattle breeds that play an essential role in the country’s agricultural economy. Breeds such as Afrikaner, Bonsmara, and Nguni are well-adapted to the region’s diverse climates, ranging from semi-arid areas to temperate zones. However, these breeds are also vulnerable to a variety of vector-borne diseases, including those transmitted by ticks and mosquitoes, which are prevalent in warmer climates (Rogers and Randolph, 2006). There were common pathogens among the South African breeds such as *Theileria parva strain muguga, Babesia bigemina & bovis, Mycoplasma* species, and *Anaplasma centrale & marginale* including strains *St. Maries, Jaboticabal*, and *Palmeira*. As with the Ugandan case, South African cattle exhibit a range of tick-borne pathogens here including *theleria, babesia, anaplasma* and *ehrlichia*. The theleria strain seen here is commonly vaccine related much like in the Ugandan case. Both *Babesia bigemina* and *bovis* have been diagnosed in south African cattle in multiple provinces and are known to cause anemia, fever and hemoglobinuria (Mtshali and Mtshali, 2013). The Anaplasma species here include *Marginale* of strains *Dawn, St Maries* and *Florida*, as well as *Ovis* of strain *Haibei*.

The analysis of South African cattle breeds—Afrikaner, Bonsmara, Holstein, Nguni, and Tuli—revealed distinct pathogen profiles for each breed, with unique pathogens identified in all breeds. The Afrikaner breed was found to have unique pathogens such as *Sarcocystis hjorti, Candidatus Mycoplasma erythrocervae*, and *Theileria parva parva. Sarcocystis* species are protozoan parasites known to affect muscle tissues, while Theileria parva is associated with East Coast fever, a tick-borne disease that can be lethal for cattle (Surve et al., 2023). The Bonsmara breed exhibited several unique pathogens, including *Candidatus Anaplasma turritanum, Arenicola marina, Anaplasma sp. NS108*, and *Trypanosoma congolense IL3000*. These pathogens are associated with diseases like anaplasmosis and trypanosomiasis, both of which can cause significant cattle morbidity and mortality due to blood-borne infections (Paoletta et al., 2018). The Holstein breed was found to harbor unique pathogens such as *Anaplasma marginale str. Washington Okanogan, Babesia sp. Xinjiang-2005, and Mycoplasma sp. China-1. Anaplasma marginale* is a well-known pathogen responsible for anaplasmosis, while Babesia species are associated with babesiosis, a parasitic disease that affects red blood cells (Paoletta et al., 2018). The Nguni breed displayed unique pathogens including *Brucella pseudintermedia, Mycoplasma putrefaciens, Anaplasma phagocytophilum str. HZ2*, and *Brucella anthropi*. These pathogens suggest potential exposure to brucellosis and anaplasmosis, both of which can have significant health implications for cattle. Lastly, the Tuli breed demonstrated a distinct pathogen profile with unique pathogens such as *Neospora caninum Liverpool, Pseudomonas aeruginosa SJTD-1, Corynebacterium bovis DSM 20582, Trypanosoma theileri*, and *Klebsiella oxytoca. Neospora caninum* is particularly concerning as it is associated with reproductive issues and abortions in cattle (Haddad et al., 2005), while Trypanosoma species can lead to trypanosomiasis, a major cattle disease in Africa (Paoletta et al., 2018; Rascón-García et al., 2023).

## Discussion

This is the most comprehensive study to date investigating the microbial diversity present in unmapped reads obtained from WGS data of cattle populations. Unmapped read data was analyzed across geographically diverse regions in Europe and Africa. These regions were selected to capture a wide range of environmental conditions and farming systems, providing a comprehensive view of how pathogen diversity may vary in response to geographic and climatic factors. We have been studying locally adapted cattle breeds in all these regions to generate novel insights into climatic and environmental adaptation of livestock, therefore these regions are bioclimatically diverse ranging from as low as –30 °C in Finland up to +50 °C in South Africa and Egypt, therefore presenting an interesting landscape to investigate pathogen presence globally. We have developed here a bioinformatics pipeline to identify pathogens in unmapped reads and filter against a custom database informed by literature searches pertinent to countries and their pathogen landscapes.

One of the most striking findings of this study is the apparent correlation with “spillover” of pathogens traditionally associated with African cattle breeds into southern European regions, particularly Portugal. Pathogens such as *Theileria parva, Anaplasma platys, Theileria orientalis*, and *Babesia bigemina*, commonly found in Uganda, South Africa, and Egypt, were also detected in Portuguese breeds. This suggests that warmer European countries, like Portugal, may be increasingly exposed to pathogens typically associated with tropical and subtropical climates. The presence of these pathogens in Portugal, a country experiencing rising temperatures, supports the hypothesis that climate change is facilitating the northward spread of vector-borne diseases. In contrast, these same pathogens were notably absent or present at much lower frequencies in northern European countries such as Finland and the Netherlands, where cooler climates likely restrict vector activity. The stark difference between pathogen profiles in northern versus southern Europe underscores the importance of climate in shaping pathogen distribution (Ainsworth, 2023). With southern Europe projected to experience even warmer and more humid conditions in the coming decades, the risk of pathogen spillover is likely to intensify, posing a growing threat to livestock health in the region.

The study revealed considerable variation in pathogen diversity across different cattle breeds. Notably, breeds from tropical and subtropical regions (e.g., Ugandan and South African breeds) showed higher pathogen alignments compared to those from temperate regions. This aligns with the well-established relationship between warmer climates and the proliferation of disease vectors such as ticks and mosquitoes. For example, Ugandan breeds such as Nkedi and Nganda demonstrated a high diversity of pathogen alignments, including Trypanosoma species and Brucella, which are known to thrive in warm, vector-rich environments.

In contrast, Finnish breeds such as Eastern and Northern Finncattle exhibited far lower pathogen alignment diversity. This may reflect not only the cooler climate but also the small-scale, biosecure farming practices typical of northern Europe. However, it is notable that certain pathogens, such as *Mycoplasma haemominutum* and *Mycoplasma ovis*, were consistently found across all Finnish breeds, indicating that some pathogens are well-adapted to colder climates. What is most peculiar in Finnish breeds is the presence of various avian coccidiosis causing pathogens. This may be due to environmental exposure. This could very well be the case with other breeds and pathogens, especially in Dutch breeds that exhibited multiple *E. coli* and salmonella species.

Nevertheless, this study is limited by its initial design. The filtered sequences attributed to pathogens are the result of a BLAST alignment, which compares nucleotide to that which is available in a database and attributes it to the statistical likelihood of this match. This by no means is a reliable indicator of pathogen presence in this case. In addition, the study is limited by the literature search, which is biased in its own way. By stringed filtering of the output, we expect to reduce the false positive results. It was found that it was much more straightforward to search for comprehensive literature about endemic disease of Ugandan cattle breeds than it was for their Finnish counterparts, for example. In addition, the k-mer analysis is based on a design for the search of viral genomes, and failed to produce a high degree of clustering. This means that while visually there are some patterns, statistically their presence cannot be confirmed. K means clustering and ARI scores have highlighted this unfortunate trend in further investigations. It is also very important to note that the sample collection also could have not been free from contamination. The pathogens found could very well not be associated with the individual sampled and could be associated with the environment of the animal and validation of the results in bloodserum samples of these animals for the specific pathogens will improve benchmarking of this approach. What this study offers is correlation to real world events and the potential development and use of a new approach in identification of the disease history of an animal.

## Supplementary data related to this article

**Supplementary file 1- Peer-reviewed research used for pathogen database**

**Supplementary file 2- Contig assembly statistics**

**Supplementary file 3- Breed specific pathogen sequence alignments**

## CRediT authorship contribution statement

**Daniil Ruuvinskiy:** Formal analysis, Methodology, Writing – original draft. ..... Writing – review & editing, **Juha Kantanen:** Supervision, Writing – review & editing. **Richard Crooijmans:** Funding acquisition, Writing – review & editing. **Kisun Pokharel:** Supervision, Writing – review & editing.

## Declaration of competing interest

The authors have no conflicts of interest to declare.

## Acknowledgements

The work was supported by European Union’s Horizon 2020 research and innovation program (grant agreement No. 727715). The authors wish to acknowledge CSC – IT Center for Science, Finland, for computational resources. Finnish Cultural Foundation is acknowledged for providing PhD grant to DR.

## Data availability

Unmapped sequences resulting from the whole genome sequence data of samples included in this study will be made available through the European Nucleotide Archive (ENA) at https://ebi.ac.uk/ena.

## Notes

https://figshare.com/articles/figure/Kronatools_charts_of_Blast_analyses_of_Unmapped_Reads/28730078

### Competing Interest Statement

The authors have declared no competing interest.

